# High proportion of multiple copies of *Plasmodium falciparum Plasmepsin-2* gene in African isolates: Is piperaquine resistance emerging in Africa?

**DOI:** 10.1101/361204

**Authors:** D. Leroy, F. Macintyre, M. Adamy, B. Laurijssens, R. Klopper, N. Khim, E Legrand, T.N. Wells, D. Ménard

## Abstract

Emergence of *Plasmodium falciparum* resistance to antimalarial drugs is currently the primary rationale supporting the development of new and well-tolerated drugs. In 2014-2015, a phase 2b clinical study was conducted to evaluate the efficacy of a single oral dose of Artefenomel (OZ439)-piperaquine (PPQ) in Asian and African patients presenting with uncomplicated *falciparum* malaria. Blood samples collected before treatment offered the opportunity to investigate the proportion of multidrug resistant parasite genotypes including *P. falciparum Kelch13* mutations and copy number variation of both *P. falciparum plasmepsin2* (*Pfpm2*) and *P. falciparum multidrug resistance 1 (Pfmdr1)* genes. Validated *Kelch13* resistance mutations including C580Y, I543T, P553L and V568G were only detected in parasites from Vietnamese patients. In Africa, isolates with multiple copies of the *Pfmdr1* gene were shown to be more frequent than previously reported (21.1%, range from 12.4% in Burkina Faso to 27.4% in Uganda). More strikingly, high proportions of isolates with multiple copies of the *Pfpm2* gene, associated to PPQ resistance, were frequently observed in the African sites, especially in Burkina Faso and Uganda (>30%).

Our findings sharply contrast with the recent description of increased sensitivity to PPQ of Ugandan parasite isolates. This emphasizes the necessity to decipher the genetic background associated with PPQ resistance in Africa by investigating *in vitro* susceptibilities to PPQ of isolates with multiple copies of the *Pfpm2 gene* and the urgent need to assess the risk of development of PPQ resistance, along with the efficacy of both current frontline therapies and new antimalarial combinations.

## Introduction

Emergence of *Plasmodium falciparum* resistance to antimalarial drugs is currently the primary rationale supporting the development of new and well-tolerated drugs. While the estimated number of malaria cases in the world decreased from 237 million (218–278 million) in 2010 to 211 million (192–257 million) in 2015, the morbidity and the mortality have stabilized in 2016 with estimates of 216 million cases (196–263 million) and 445,000 deaths (compared to 446,000 in 2015) as reported by the WHO (1-3). Globally, the vast majority of deaths (>90%) caused by malaria is due to *P. falciparum* infections occurring in Africa, children under five years of age. Artemisinin Combination Therapies (ACTs) which are currently recommended as first-line treatment of uncomplicated *falciparum* malaria, are less effective in Southeast Asia, particularly in Cambodia, where high rates of treatment failure associated with artemisinin and piperaquine resistance are currently reported (4-16). The containment and the elimination of these multidrug resistant parasites in Southeast Asia are a priority for the WHO to avoid their spread to Africa as was the case with previous generations of antimalarial drugs (e.g. chloroquine, sulfadoxine-pyrimethamine) (17). Fortunately, molecular markers associated with such resistance are available (10). In particular, mutations in the propeller domain of a *Kelch* gene located on the chromosome 13 (*Kelch13*), and amplification of a cluster of genes encoding both *Plasmepsin 2* (*Pfpm2*) and *Plasmepsin 3* proteins, have been recently shown to be associated with artemisinin and PPQ resistance, respectively (18-20).

According to the latest WHO update on artemisinin resistance (21), to be validated a *Kelch13* resistance mutant has to be correlated with delayed parasite clearance in clinical studies and reduced drug *in vitro* susceptibility (survival rate ≥ 1% expressed by the Ring-stage Survival Assay, RSA0-3h) in fresh isolates (*ex vivo* assays), or culture-adapted field parasites or *Kelch13* genome-edited parasites (*in vitro* assays) (22-25). To date, only five *Kelch13* mutations are validated (C580Y, Y493H, R539T, I543T, N458Y). The F446I mutant, which is highly prevalent in Myanmar, is strongly suspected of being associated to artemisinin resistance. In Africa, a broad array of rare non-synonymous mutations in the *Kelch13* gene have been described in *P. falciparum* isolates, but any of these mutants have not been associated with artemisinin resistance (26), attesting that not all non-synonymous *Kelch13* mutations confer resistance to artemisinin.

More recently, resistance to PPQ has been associated with an increase of survival rates of parasite exposed to 200 nM PPQ for 48 hours (piperaquine survival assay, PSA) and with the amplification of *plasmepsin 2-3* genes (*Pfpm2-3*) (6, 20). In Cambodia, where high rates of treatment failure to dihydroartemisininpiperaquine (DHA-PPQ) are observed (i.e. >60% in some provinces), it has been demonstrated that amplification of *Pfpm2* gene and presence of validated *Kelch13* mutations were highly predictive of DHA-PPQ treatment failure (20). Most of these parasites harbor a single copy of *Pfmdr1* gene leading to the recovery of mefloquine sensitivity (4, 6) and suggesting a natural antagonism between PPQ resistance and mefloquine resistance. However, we still do not understand whether *Pfmdr1* de-amplification (from multiple copies to single copy *Pfmdr1*) is due to the implementation of DHA-PPQ as first-line treatment or due to the release of mefloquine pressure and an increase in parasite fitness accompanying *Pfmdr1* gene de-amplification. To date, DHA-PPQ resistance is confined to Southeast Asia. Only a recent study conducted in Mozambique has provided evidence of the presence (at a very low frequency, 1.1%) of parasites carrying multiple copies of *Pfpm2* (27).

Facing the threat of losing all current ACTs front-line therapies due to resistance, new generation of endoperoxides with more favorable pharmacokinetic profiles like the ozonide Artefenomel (OZ439) have been developed (28). The efficacy of this new chemical entity was recently evaluated in combination with PPQ in African and Southeast Asian (Vietnam) patients with uncomplicated *falciparum* malaria infection (29). The primary objective of this phase 2b clinical study was to determine whether a single oral dose combination of artefenomel/PPQ was an efficacious and safe treatment (e.g., ≥ 95% of patients cured on the basis of polymerase chain reaction (PCR)-adjusted Adequate Clinical Parasitological Response at Day 28 (ACPR28)) for adults and children infected by *P. falciparum*. Blood samples collected in 2014-2015 from this clinical trial offered the opportunity to investigate the proportion of multidrug resistant parasites (*i.e. P. falciparum Kelch13* mutants and gene copy number of both *Pfpm2* and *Pfmdr1*). Here, we report the occurrence of such genotypes from these samples and provide a map of potential risk of emergence of resistance to the main front-line therapies currently used to treat malaria-infected patients and to the next generation of antimalarial combinations.

## Results

The *P. falciparum* samples collected from patients before treatment and yielding a successful result, by country and molecular assay, are presented in ***Table 1***.

**Table 1.**
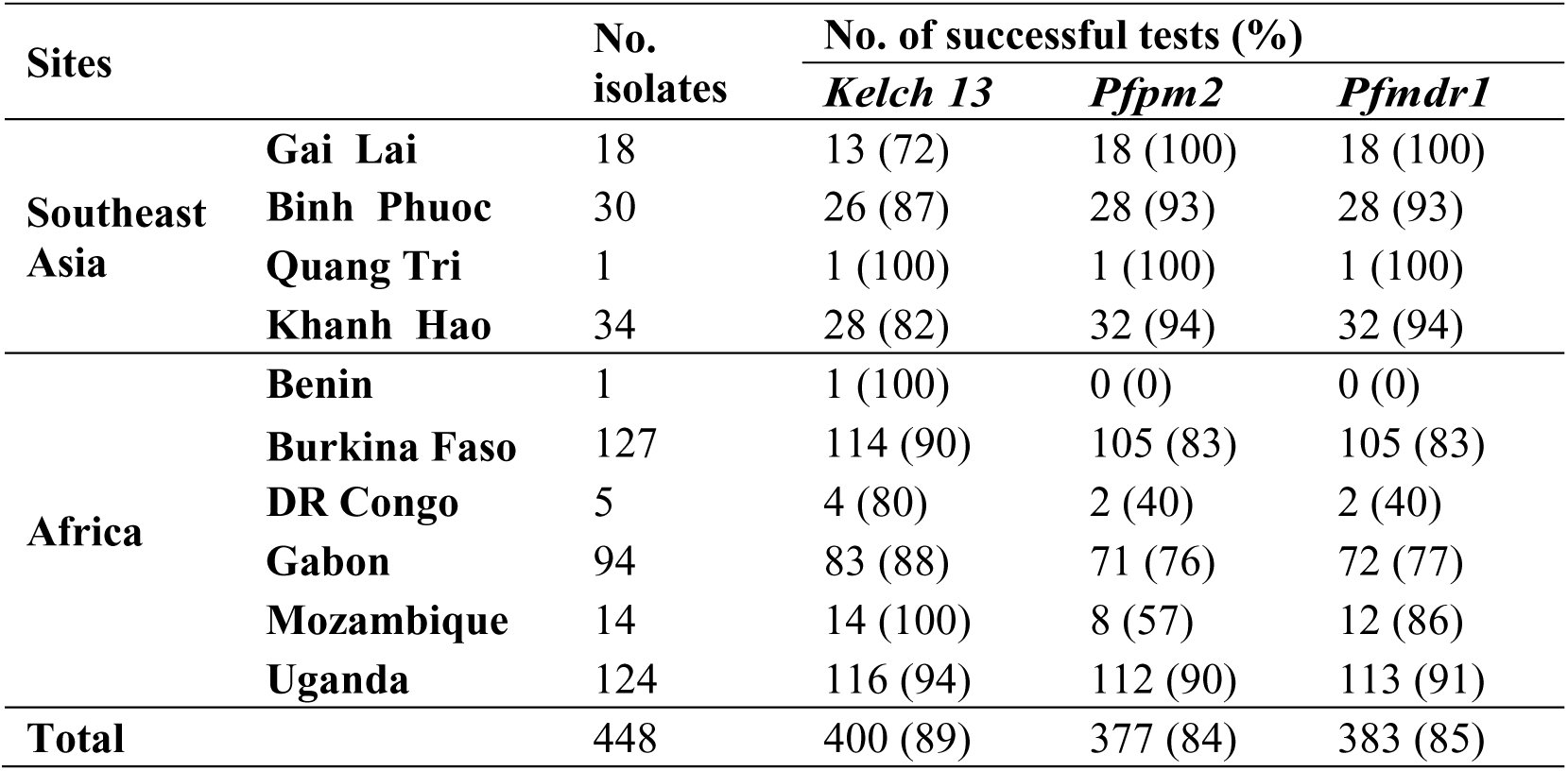
Number of isolates collected from each sites in Southeast Asia (Vietnam) and Africa and number and proportion of successful molecular tests.

### Global genotypes overview

Among the 68 Southeast Asian clinical isolates collected in Vietnam with available data, 67.6% (46/68) were found to harbor parasite with validated or candidate *Kelch13* resistance mutations (***Table 2***). Details regarding *Kelch13* mutants according to the collection sites are presented in ***Table 3***. In contrast, none of the 332 isolates collected from African patients and successfully tested were found to carry validated or candidate *Kelch13* resistance mutations.

**Table 2.** Distribution (number and proportion) of genotypes (*Kelch13* mutation*s, Pfmdr1* and *Pfpm2* gene copy numbers) detected in *Plasmodium falciparum* isolates collected from Southeast Asia and Africa in 2014-2015.

| Locus | Allele/Haplotype | Number of isolates (%) detected in |  | P-value |
| --- | --- | --- | --- | --- |
|  |  | Asia (N=82) | Africa (N=355) |  |
| *Kelch 13* | **ART** | **46 (67.6)** | **0 (0.0)** | < 10<sup>-4</sup> |
|  | OTH | 1 (1.5) | 10 (3.0) |  |
|  | WT | 21 (30.9) | 322 (97.0) |  |
| *Pfpm2* | Single copy | 68 (86.1) | 218 (73.2) | 0.02 |
|  | **Multiple copies** | **11 (13.9)** | **80 (26.8)** |  |
| *Pfmdr1* | Single copy | 74 (93.7) | 240 (78.9) | 0.002 |
|  | Multiple copies | 5 (6.3) | 64 (21.1) |  |
| *Pfpm2/Pfmdr1* | Single copy/Single copy | 64 (81.0) | 189 (63.4) | 0.009 |
|  | Single copy/Multiple copies | 4 (5.1) | 29 (9.7) |  |
|  | **Multiple copies/Single copy** | **10 (12.7)** | **47 (15.8)** |  |
|  | Multiple copies/Multiple copies | 1 (1.3) | 33 (11.1) |  |
| *Kelch 13/Pfpm2/Pfmdr1* | **ART/Multiple copies/Single copy** | **7 (10.6)** | **0 (0.0)** | < 10<sup>-4</sup> |
|  | WT/Multiple copies/Single copy | 2 (3.0) | 43 (14.7) |  |
|  | ART/others | 38 (57.5) | 0 (0.0) |  |
|  | WT/others | 18 (27.3) | 241 (82.5) |  |

ART: validated or candidate *Kelch 13* mutations; WT: *Kelch 13* Wild type, OTH: *Kelch 13* mutations with unknown association with artemisinin resistance (detailed in Tables 3 and 4). Bold font denotes the allele or haplotype associate with drug resistance. P-value (Chi-squared test<sup>±</sup> or Fischer exact test<sup>¥</sup>)

**Table 3.**
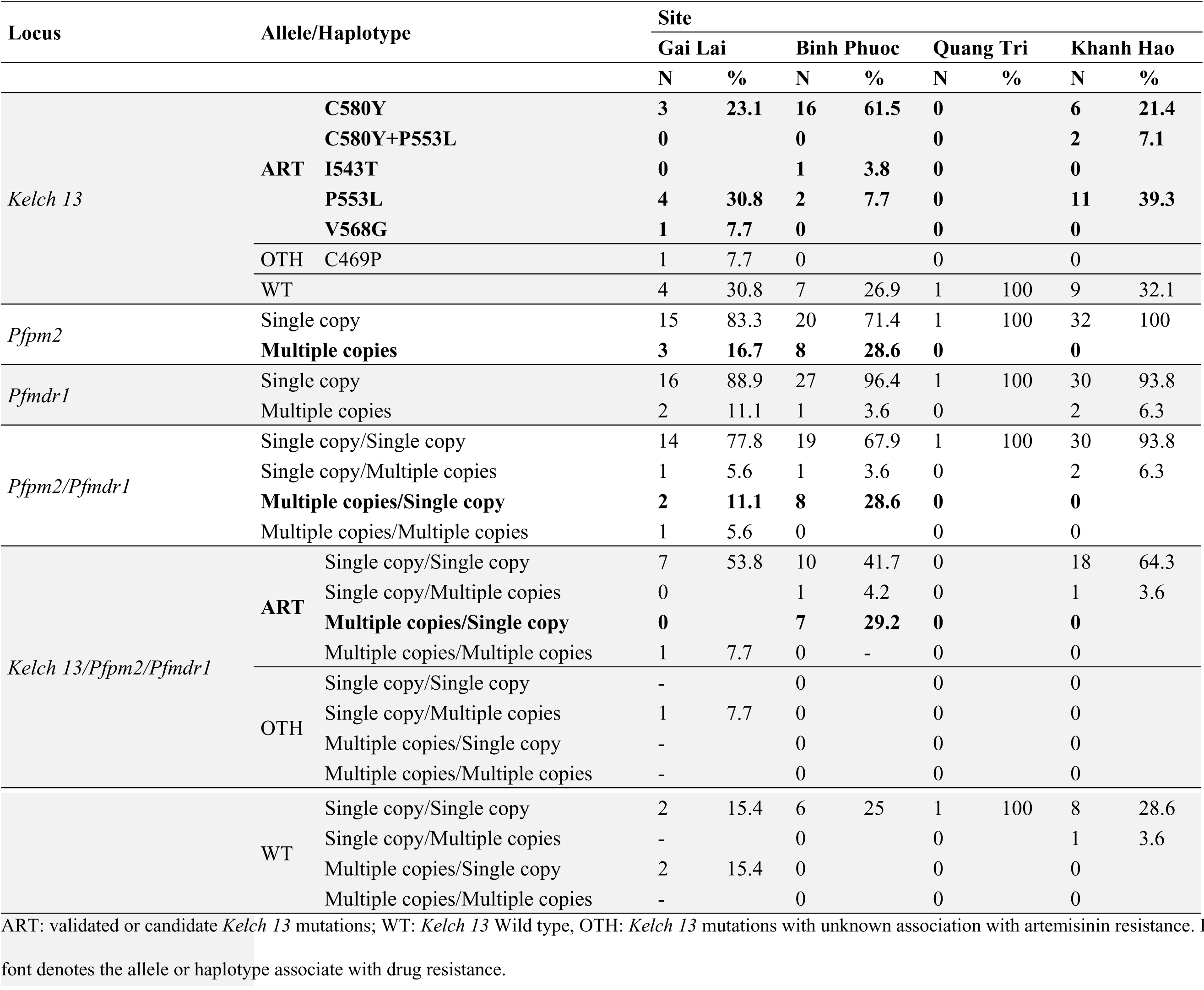
Distribution (number and proportion) of genotypes (*Kelch13* mutation*s, Pfmdr1* and *Pfpm2* gene copy numbers) detected in *Plasmodium falciparum* isolates collected in four sites located in Southeast Asia in 2014-2015.

Significant difference in proportion of isolates with multiple copies *Pfmdr1* were found between Africa (21.1%, 64/304, 95%CI:16.2-26.9%) and Asia (6.3%, 5/79, 95%CI:2.0-14.8%, p=0.002, ***Table 2***). Parasites with multiple copies of *Pfpm2* were observed in 11 Asian samples (13.9%, 11/79, 95%CI:6.9-24.9%) and unexpectedly at higher proportion in African isolates (26.8%, 80/298, 95%CI:21.3-33.4%, p=0.02, ***Table 2***). However, multiple copies of *Pfpm2*/single copy *Pfmdr1*, hypothesized to favour resistance to PPQ, were found at similar proportion in 10 Asian isolates (12.7%, 10/79, 95%CI:6.1-23.3%) and 47 African samples (15.8%, 47/298, 95%CI:11.6-21.0%, p=0.72, ***Table 2*** and ***Figure 1***).

**Figure 1.**
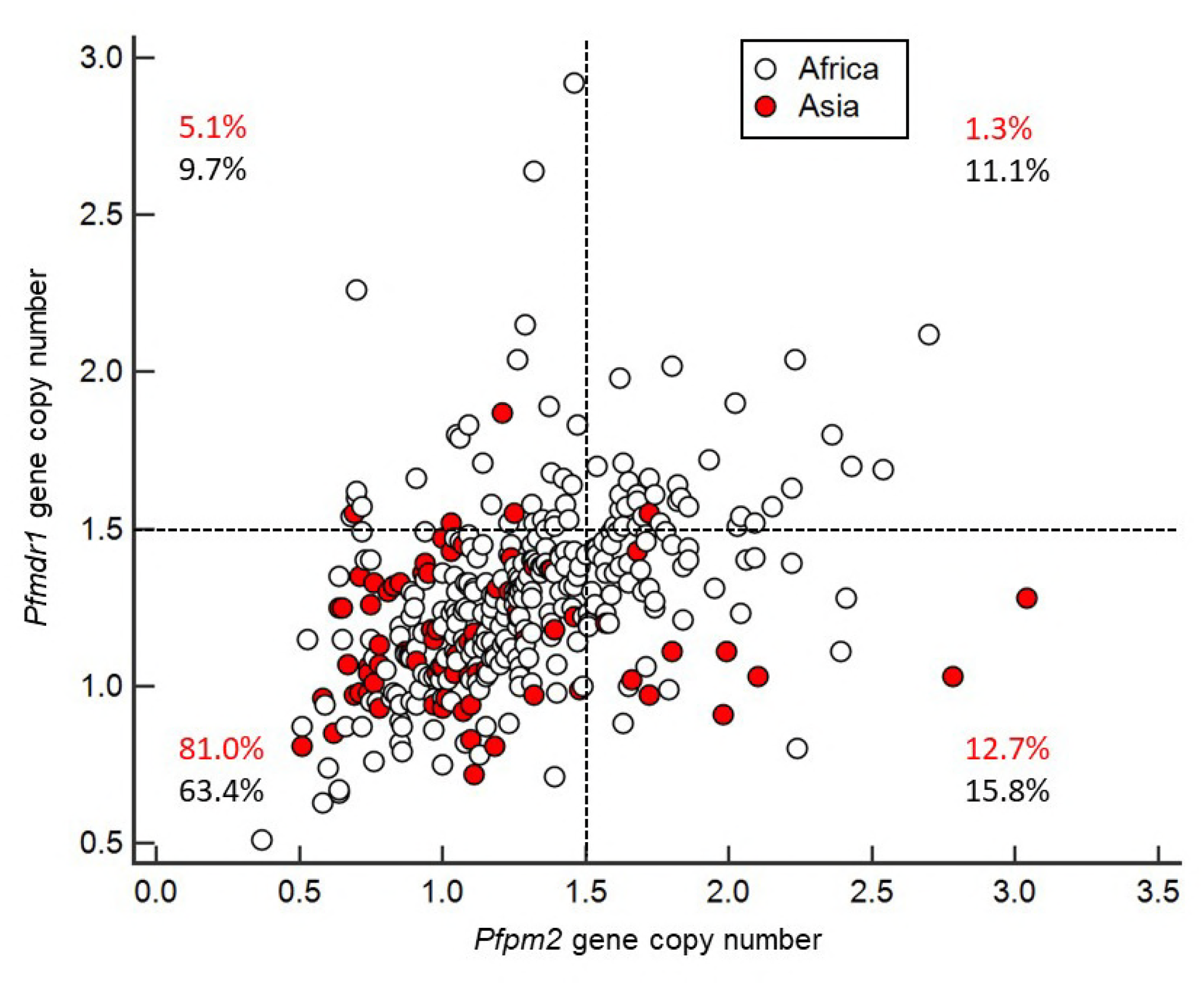
*Pfmdr1* and *Pfpm2* gene copy numbers of *Plasmodium falciparum* isolates collected from Southeast Asia (in red) and from Africa (in black). Proportion of the isolates from Southeast Asia and African are given for each group: *Pfpm2* single copy/*Pfmdr1* single copy (lower-left quandrant), *Pfpm2* single copy/*Pfmdr1* multiple copies (upper-left), *Pfpm2* multiple copies/*Pfmdr1* single copy (lower-right) and *Pfpm2* multiple copies/*Pfmdr1* multiple copies (upper-right).

In Asia, seven isolates (10.6%, 7/65, 95%CI:4.3-22.2%) had genotypes associated with both artemisinin and PPQ resistance (i.e. with *Kelch13* validated and candidate resistance mutations, and multiple copy *Pfpm2*/single copy *Pfmdr1*) (***Figure 2, panel A***). In Africa, no clinical isolates had mutations conferring both artemisinin and PPQ resistance due to the absence of *Kelch13* mutant-type parasites (***Figure 2, panel B***).

**Figure 2.**
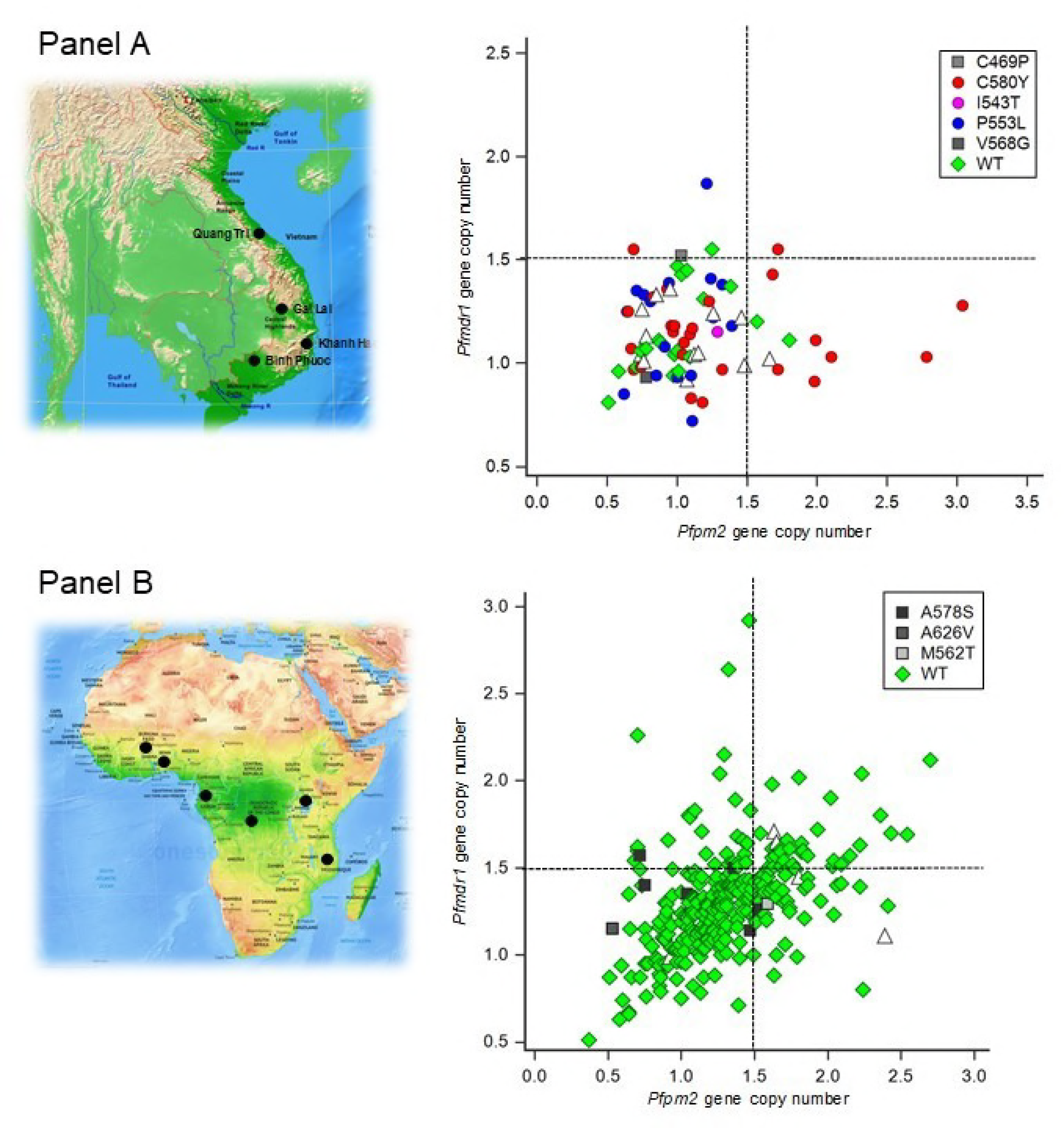
*Kelch13* mutation*s, Pfmdr1* and *Pfpm2* gene copy numbers of *Plasmodium falciparum* isolates collected from 4 sites in Southeast Asia (Panel A) and from 9 sites in Africa (Panel B). Each *Kelch13* mutation*s* are presented with different symbols and colours. Open triangle represents isolates with unavailable *Kelch13* data. The four quadrants in both panel presents isolates with *Pfpm2* single copy/*Pfmdr1* single copy (lower-left quandrant), *Pfpm2* single copy/*Pfmdr1* multiple copies (upper-left), *Pfpm2* multiple copies/*Pfmdr1* single copy (lower-right) and *Pfpm2* multiple copies/*Pfmdr1* multiple copies (upper-right).

### Southeast Asian (Vietnamese) genotypes (Table 3)

*Kelch13* validated and candidate mutations were detected in >60% of the isolates in all sites (from 61.1% in Gai Lai to 73.0% in Binh Phuoc) except Quang Tri (where only one sample was collected). C580Y was the most predominant *Kelch13* validated and candidate mutation (54.3%, 25/46, 95%CI:25.2-80.2%) followed by P553L (37.0%, 17/46, 95%CI:21.5-59.2%), I543T (2.2%, 1/46, 95%CI:0.5-12.1%) and G568G (2.2%, 1/46, 95%CI:0.5-12.1%). In Khanh Hao, two isolates were found to have both C580Y and P553L single mutant parasites (likely from polyclonal infections).

Isolates with multiple copies of *Pfpm2* were detected only in two sites located along the Cambodian border: in Gai Lai (16.7%, 3/18, 95%CI:3.4-48.7%) and in Binh Phuoc (28.6%, 8/28, 95%CI:12.3-56.3%). No parasites with multiple copies were detected out of 32 isolates in Khanh Hao. Parasites with a single copy of *Pfmdr1* were frequent (>88%) in samples collected from all four study sites (from 88.9% in Gai Lai to 100% in Quang Tri).

Parasites with multiple copies *Pfpm2*/single copy *Pfmdr1* were observed in 10/79 (12.6%, 95%CI:6.1-23.3%) of the isolates collected from Vietnamese patients, representing in Gai Lai (11.1%, 2/18, 95%CI:1.4-40.1%) and in Binh Phuoc (28.6%, 8/28, 95%CI:12.3-56.3%).

Isolates with genotype conferring both artemisinin and PPQ resistance (i.e. with *Kelch13* validated and candidate mutations, and multiple copy *Pfpm2*/single copy *Pfmdr1*) were only observed in patients enrolled in Binh Phuoc (29.2%, 7/24, 95%CI:11.7-60.0%).

### African genotypes (Table 4)

**Table 4.**
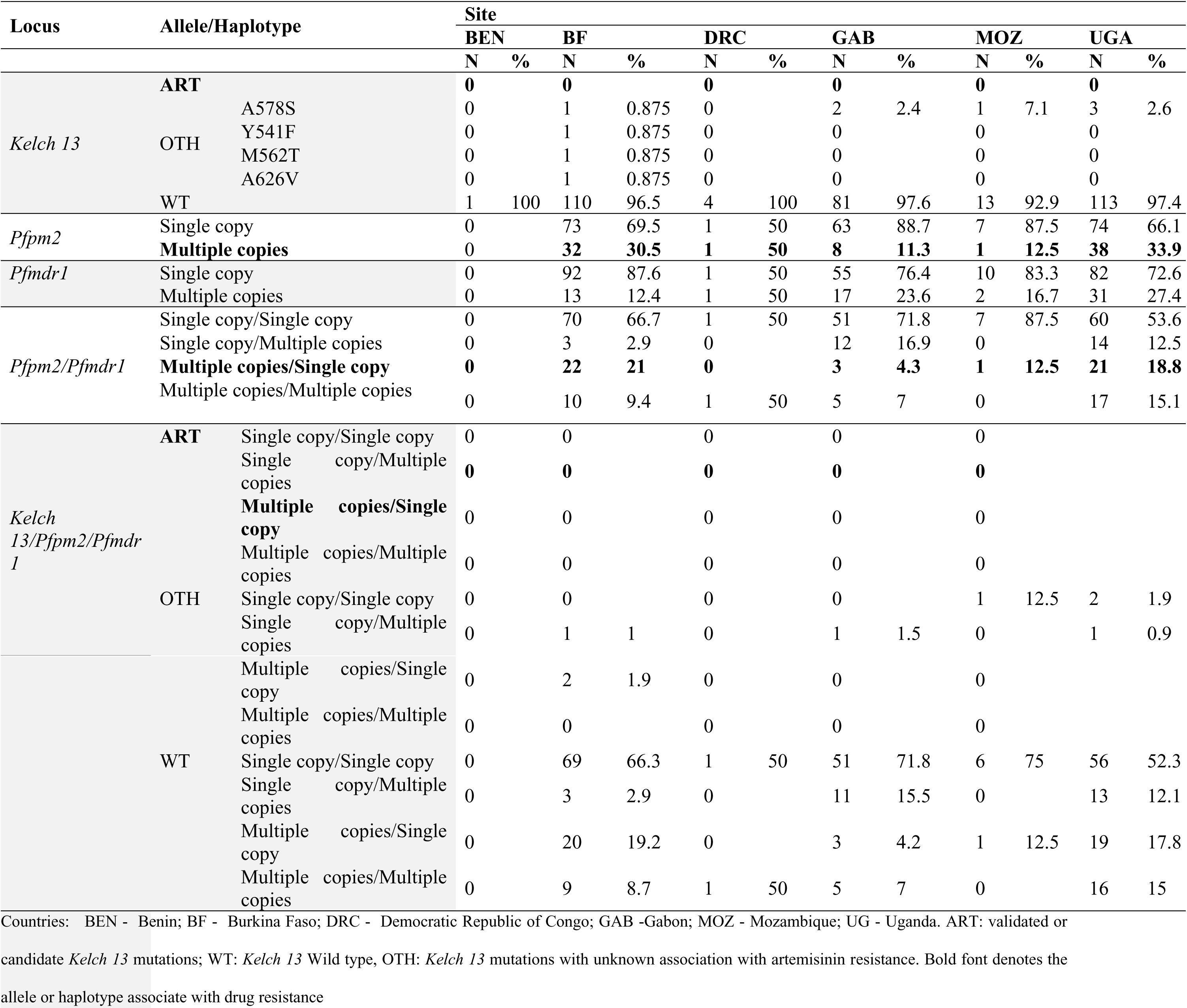
Distribution (number and proportion) of genotypes (*Kelch13* mutation*s, Pfmdr1* and *Pfpm2* gene copy numbers) detected in *Plasmodium falciparum* isolates collected in nine sites located in Africa in 2014-2015.

No *Kelch13* validated and candidate mutations were detected at any site. Other non-synonymous mutations were observed: A578S was the most predominant *Kelch13* mutation (7/10; 3 in Uganda, 2 in Gabon, 1 in Mozambique and 1 in Burkina Faso) followed by Y541F, M562T and A626V (only detected once in isolates from Burkina Faso).

Isolates from Uganda and Burkina Faso showed an unexpected high frequency of parasites with multiple copies of *Pfpm2* (34.0%, 38/112, 95%CI:24.0-46.6% and 30.5%, 32/105, 95%CI:20.9-43.0%, respectively). Samples from Gabon and Mozambique had a lower frequency of multiple copies of *Pfpm2* estimated at 11.3% (8/71, 95%CI:4.9-22.2%) and 12.5% (1/8, 95%CI:0.3-69.6%) respectively. Of note, in the Democratic Republic of Congo, results from two isolates were available and one isolate was found to carrying parasites with multiple copies of *Pfpm2*.

Parasites with single copy *Pfmdr1* were detected in almost all isolates in patients enrolled across the six African sites, therefore only 13/105 (12.4%, 95%CI:6.6-21.2%) isolates from Burkina Faso, 2/12 (16.7%, 95%CI:2.0-60.2%) from Mozambique, 17/72 (23.6%, 95%CI:13.8-37.8%) from Gabon and 31/113 (27.4%, 95%CI:18.6-38.9%) from Uganda had multiple copies of *Pfmdr1*. One out of two patients harbored parasites with multiple copies of *Pfmdr1* in DRC.

Parasites with multiple copies *Pfpm2*/single copy *Pfmdr1* were observed at a frequency of 21.0% (22/105, 95%CI:13.1-31.7%) in Burkina Faso, 18.8% (21/112, 95%CI:11.6-28.7%) in Uganda, 12.5% (1/8, 95%CI:0.3-69.7%) in Mozambique and 4.3% (3/71, 95%CI:0.9-12.5%) in Gabon. However, isolates with genotype conferring both artemisinin and PPQ resistance (i.e. with *Kelch13* validated and candidate mutations, and multiple copy *Pfpm2*/single copy *Pfmdr1*) were not observed in patients enrolled in Africa since there were no *Kelch13* validated and candidate mutations.

## Discussion

The current phase 2b clinical study of Artefenomel, an ozonide showing improved pharmacokinetics properties compared to artemisinins, combined with PPQ was designed to assess the efficacy of single oral doses in patients with uncomplicated falciparum malaria in Southeast Asia (Vietnam) and Africa. In addition to the clinical outcome assessment, we investigated in isolates collected before treatment, three molecular markers associated with drug resistance for mapping the potential risks of future treatment failures. The frequency of *Kelch13* mutations associated with artemisinin resistance, and *Pfmdr1* and *Pfpm2* genes copy number were measured in available isolates collected from all clinical sites. Our investigations confirmed that artemisinin resistance is still confined in Southeast Asia. We observed a high proportion of *Kelch13* validated and candidate resistance mutations as well as a new unreported one (C469P) in Vietnamese parasites and the complete absence of these mutants in African isolates. As previously reported (26, 30, 31), we detected in our African samples a low proportion of *Kelch13* mutations and all these mutations have not been shown to be associated to artemisinin resistance (26).

However, we observed a higher proportion (3-fold) of parasites with multiple copies of *Pfmdr1*, a gene encoding a drug efflux pump, in African samples compared to Southeast Asian isolates. This observation contrasts with previous reports showing high frequency of parasites with multiple copies of *Pfmdr1* in Asia (32-34) compared to Africa (35-37). These finding likely reflect the profiles of evolution of *P. falciparum* populations linked to antimalarial drug pressure in both continents. Especially, the prevalence of high *Pfmdr1* amplification observed in Africa might be linked with the routine use of artemether-lumefantrine as first line treatment for more than a decade. Indeed, increased *pfmdr1* copy number is known to modulate parasite responses to a wide range of drugs including lumefantrine (35, 38, 39). Supporting this expectation, it seems feasible that such parasites exposed to lumefantrine as monotherapy for several days following clearance of artemether have been selected, while parasites with a single copy have been eliminated. In contrast, the low prevalence of *Pfmdr1* multiple copies observed in Southeast Asia could be due to the recent implementation of DHA-PPQ, the removal of the mefloquine drug pressure or both, as the case in Cambodia (18, 20, 40).

High frequency of isolates with multiple copies of the *Pfpm2* has already been reported in recent studies conducted in Cambodia (18, 20, 40). As the Vietnamese clinical sites (Gai Lai and Binh Phuoc) are located alongside the Cambodian border (***Figure 2***), we can suspect that data from our study might reflected an evolving situation where the amplification of *Pfpm2* is spreading beyond Cambodia, as described recently (5, 7). To date, frequencies observed in Vietnamese isolates are not yet as high as the ones observed in Cambodia but might continue to increase in the future.

Unexpectedly, in African isolates, amplification of *Pfpm2* gene was shown to occur at a much higher frequency (~27% on average across clinical sites in Africa, reaching 30.5% in Burkina Faso and 33.9% in Uganda) than was recently described (from 11.1% to 13.8% in Uganda and 1.1% in Mozambique) (27, 41). Considering the geographical extent and the diversity of the clinical sites in Africa, the high frequency reported at sites distant to each other suggests that amplification of *Pfpm2* gene occurred independently in each site. More importantly, since in Southeast Asia most parasites with multiple copies of *Pfpm2* also display *Kelch13* resistance mutations, which is not the case in African samples, it is likely that *Pfpm2* amplification originated in Africa, independently of Southeast Asia.

Unfortunately, we were not be able to perform *in vitro or ex-vivo* drug susceptibility assays and test association between *Pfpm2* amplification and clinical resistance to PPQ in the current study. An evaluation is currently ongoing to see whether, and if so to what extent, these markers of artemisinin and PPQ resistance affected the parasite clearance half-life (PCT1/2) and PCR-adjusted 28 days follow up in patients treated with artefenomel/PPQ (study MMV OZ439 13 003). However, it was recently reported that compared to drug sensitivities measured on Ugandan isolates from 2010 to 2013 (from the same site, namely Tororo), those measured in 2016 to chloroquine, amodiaquine, and PPQ were increased by 7.4, 5.2 and 2.5-fold respectively (41). This longitudinal study showed that rather than drug resistance developing to these three antimalarial drugs, an increase in sensitivity was observed that was correlated with low prevalence of the polymorphisms recently associated with resistance to artemisinins or PPQ. Indeed, clinical resistance to DHA-PPQ has not yet been reported in Africa (42). Although, we cannot exclude the possibility that parasites showing amplification of *Pfpm2* observed in the current study are resistant to PPQ without confirmation of *in vitro* or *ex vivo* phenotypes, data reported by Rasmussen et *al*. (41) suggest that significant occurrence of clinical resistance to PPQ is unlikely. In other words, in Africa it is unclear whether the amplification of *Pfpm2* is necessary and/or sufficient for the development of resistance to PPQ. The ongoing analysis relating the markers of resistance to clinical outcome may provide some insights regarding this question. It is still debated whether additional genetic modifications in the *P. falciparum chloroquine resistance transporter* gene are required to confer such resistance (43, 44). Indeed, recent genomic and biological investigations have revealed a rapid increase in the prevalence of novel *Pfcrt* mutations in Cambodia (H97Y, F145I, M343L, and G353V). These mutants (from culture-adapted Cambodian field isolates or Dd2 gene-edited clones) were confirmed to confer PPQ resistance as determined using the PSA^0-3h^ (6, Ross et al. in revision).

At present, several ACTs are used in Africa and Asia to treat patients with uncomplicated malaria. artemether-lumefantrine (AL), artesunate-amodiaquine (AS-AQ), artesunate-mefloquine (AS-MQ), artesunate-sulfadoxine-pyrimethamine (AS-SP), dihydroartemisinin-piperaquine (DHA-PPQ) and pyronardidine-artesunate (PA). All achieve more than 95% efficacy in clinical trials based on PCR-adjusted Day28 ACPR. Due to the long post treatment prophylaxis of the well-tolerated PPQ, DHA-PPQ is currently under evaluation in a number of interventions such as Intermittent Preventive Treatment in pregnant women or in infants (IPTp, IPTi) and Mass Drug Administration campaigns (MDA) in Africa. As a key surveillance goal, it is therefore of particular importance to continue following the evolution of *Pfpm2* amplification along with mutations in the *Pfcrt* gene and to investigate whether these genetic signatures are associated with PPQ resistance in Africa.

## Material & Methods

### Study Design, study sites and population

Study MMV OZ439 13 003 was a randomized, double-blind, single-dose study to investigate the efficacy, safety, tolerability and pharmacokinetics of Artefenomel (OZ439) 800 mg in loose combination with three doses of PPQ phosphate (640, 960, 1440 mg) in male and female patients aged ≥ 6 months to < 70 years, with uncomplicated *falciparum* malaria in Africa and Southeast Asia (Vietnam), as previously described (29). This study was conducted in 13 sites, including Burkina Faso (3 sites, N=127), Uganda (1 site, N=124), Benin (1 site, N=1), the Democratic Republic of Congo (1 site, N=5), Gabon (2 sites, N=94), Mozambique (1 site, N=14), and Vietnam (4 sites, N=83). A total of 448 patients were randomized into each of three treatment arms: OZ439 800 mg/PPQ 640 mg (N=148), OZ439 800 mg/PPQ 960 mg (N=151) and OZ439 800 mg/PPQ 1440 mg (N=149).

### DNA extraction

*P. falciparum* DNA was extracted from dried blood spots using the QIAamp DNA Mini kit (Qiagen, Germany), according to the manufacturer’s instructions. Samples were screened to confirm the presence of *P. falciparum* DNA using first a qualitative real-time PCR assay targeting the *Plasmodium cytochrome b* gene and secondly on positive samples, four real-time PCR assays specifically amplifying *P. falciparum, P. vivax, P. ovale and P. malariae* (45).

### Detection of *Kelch13* mutations

*P. falciparum* positive samples were tested for the presence of mutations in the propeller domain of the *Kelch13* gene (PF3D7_1343700) that have recently been associated with artemisinin resistance (19). Amplification of the Kelch-propeller domain (codons 440-680, 720 bp) was performed as previously described (26). Cross-contamination was evaluated by adding no template samples (dried blood spots negative for *P. falciparum*) in each PCR run. PCR products were sequenced by Macrogen (Seoul, Korea). Electropherograms were analysed on both strands, using PF3D7_1343700 as the reference sequence. The quality of the procedure was assessed by including dried blood spots with known *Kelch13* mutations (wild-type, C580Y, R539T, I543T, Y493H) which were tested blindly in the same batches (each 96-well) with the test samples. Isolates with mixed alleles were considered as mutant. Following the WHO recommendations, *Kelch13* mutants were classified in our study in three groups: wild-type group (parasites with no synonymous or non-synonymous mutations compared to 3D7 sequence), *Kelch13* validated (N458Y, Y493H, R539T, I543T, C580Y) and candidate mutation (P441L, F446I, G339A, P553L, V568G, P574L, A675V) group, and other *Kelch13* mutants group (parasites with synonymous or non-synonymous mutations not present in the *Kelch13* validated and candidate resistance mutation group).

### *Pfpm2* and *Pfmdr1* genes copy number variation assessment

*Pfpm2* (PF3D7_1408000) and *Pfmdr1* (PF3D7_0523000) genes copy number were measured by qPCR using a CFX96 real-time PCR machine (Bio-Rad, France), relative to the single copy of the β-tubulin gene (used as reference gene), as previously described (20). Amplification was carried out in triplicate. In each amplification run, six replicates using DNA from 3D7 parasite reference clone and three replicates without template (water) used as negative controls were included. Copy numbers were calculated using the formula: copy number= 2^−ΔΔCt^; with ΔΔC_t_ denoting the difference between ΔC_t_ of the unknown sample and ΔC_t_ of the reference sample (3D7). Specificities of *Pfpm2* and *Pfmdr1* amplification curves were evaluated by visualizing the melt curves. Multiple copies vs single copy, of both Pfmdr1 and PfPm2, were defined as copy numbers <1.5 and ≥1.5 respectively.

### Statistical analysis

Data were recorded and analyzed using Excel software and MedCalc (MedCalc Software, Belgium). Groups were compared using the Chi squared test or the Fisher’s exact test. All reported *P-*values are two-sided and were considered statistically significant if <0.05.

### Ethical statement

The study (MMV OZ439 13 003) conformed to the Declaration of Helsinki and Standard Operating Procedures that meet current regulatory requirements and guidelines laid down by the International Conference on Harmonization for Good Clinical Practice in Clinical Studies, and approved by the relevant Independent Ethics Committees (IEC), national Institutional Review Boards and where relevant, local regulatory authorities at each of the participating sites. The study protocol was registered and the study results are reported on clinicaltrials.gov (NCT02083380).

## Acknowledgements

We are grateful to the patients who took part in the study and their families. We would like to thanks all local investigators who have conducted the clinical studies (Yeka Adoke, Alfred B Tiono, Tran Thanh Duong, Ghyslain Mombo-Ngoma, Marielle Bouyou-Akotet, Halidou Tinto, Quique Bassat, Saadou Issifou, Afizi Kibuuka, Antoinette Kitoto Tshefu, Peter G Kremsner, Bui Quang Phuc, Alphonse Ouedraogo, Michael Ramharter) and MMV staff involved in study conduct, data collection and reporting (Helen Demarest, Stephan Duparc, Sophie Biguenet). MMV would also like to acknowledge their development partner Sanofi-Aventis.

The study was funded by Medicines for Malaria Venture (MMV). MMV is funded by a number of donors. Unrestricted funding from a number of donors including; US Aid, Bill and Melinda Gates Foundation, UK Department for International Development, Norwegian Agency for Development Cooperation, Irish Aid, Newcrest Mining Limited, Australian Aid, Swiss Agency for Development and Co-operation and Wellcome Trust, contributed to the study. Study activities at the CERMEL, Gabon were supported financially by the Federal Ministry of Science, Research and Economy of Austria as part of the EDCTP programme. These activities at the Gabonese site are part of the EDCTP2 programme activities of Austria supported by the European Union. These funders had no role in the design, conduct or analysis of the trial.

